# Inbred and outcrossed *Drosophila* embryos diverge in stress defense and developmental maintenance during heat stress

**DOI:** 10.64898/2026.10.01.754988

**Authors:** Subhashis Natua, Divisha Biswas, Faizan Rashid, Auinash Kalsotra, Anna Marie Sokac

## Abstract

Early embryos must mount stress defenses without compromising the developmental programs that drive morphogenesis, but whether genetic background alters this balance remains unclear. We compared cellularizing *Drosophila melanogaster* embryos from inbred and outcrossed backgrounds under control conditions and after acute or chronic heat-stress. Both backgrounds developed successfully and showed modest transcriptomic differences at 18°C. Heat stress at 32°C, however, sharply separated their phenotypic and transcriptional responses. Inbred embryos showed poor larval hatching after acute and chronic heat, and this deficit persisted through adult eclosion. Their transcriptomes exhibited a greater stress-response burden, with enrichment of proteostasis, DNA repair, mitochondrial maintenance, RNA processing, and metabolic remodeling programs. Outcrossed embryos retained substantially greater developmental survival and higher abundance of transcripts associated with development, adhesion, epithelial organization, signaling, transport, sensory responses, and morphogenesis. Background-dependent divergence was greatest during acute heat. Under chronic heat, background-dependent differences were fewer, with most overlapping those seen during acute heat and a smaller set detected only under chronic heat. Transcript-origin analysis further showed that acute outcrossed-biased genes were predominantly maternally derived and showed no enrichment for purely zygotic transcripts, whereas chronic outcrossed-biased genes contained a larger zygotic component. In summary, the less viable inbred background mounted the broader molecular stress signature, whereas the more viable outcrossed background retained greater abundance of development-associated transcripts. These findings identify genetic background as a major determinant of how embryonic heat responses are distributed between molecular stress defense and developmental gene expression.

## Introduction

Early embryogenesis unfolds through a tightly coordinated sequence of cell divisions, cell-fate specification, tissue rearrangements, and morphogenetic events (Farrell and O’Farrell 2014; Wieschaus 2016; Stathopoulos and Newcomb 2020; Sokac et al. 2023). Heat stress challenges this coordination at several levels, disrupting protein folding, organelle function, transcription, and cytoskeletal organization (Graziosi et al. 1983; Walter et al. 1990; Figard et al. 2019; Gibbs et al. 2025; Biel et al. 2026). Although activation of the heat-shock response can limit molecular damage, the earliest *Drosophila* embryos are developmentally constrained in their ability to mount this response. Heat-shock factor remains cytoplasmic until defined nuclear cycles, delaying the capacity to induce Hsp70 during early embryogenesis (Wang and Lindquist 1998). Maternal provisioning can partially offset this vulnerability, and increasing maternal loading of the small heat-shock protein Hsp23 improves embryonic thermal tolerance, influencing subsequent developmental performance (Lockwood et al. 2017). Embryonic survival under heat may therefore depend not only on the strength of canonical stress-pathway activation, but also on whether embryos can preserve the developmental programs already in progress.

Thermal tolerance varies substantially among *Drosophila* genetic backgrounds and natural populations. Embryos derived from tropical populations tolerate acute heat better than those from temperate populations, with this divergence associated in part with differences in constitutive and inducible oxidative-stress programs (Mikucki et al. 2024). Temperature can also reveal regulatory variation that is comparatively muted under permissive conditions. For example, Oregon-R and Samarkand strains showed relatively limited expression divergence at 18°C but extensive genotype-dependent expression differences at temperature extremes, consistent with environmental stress exposing previously buffered regulatory variation (Chen et al. 2015). These studies establish that genotype influences thermal tolerance and gene-expression plasticity. However, it remains unclear how genetic background shapes the balance between molecular stress defense and continued developmental gene expression in the early embryo.

Recent work using this embryonic heat-stress paradigm suggests that this balance is biologically consequential. Heat exposure induces heat-shock and endoplasmic-reticulum stress responses and broadly perturbs developmental and other cellular gene-expression programs (Biel et al. 2026; Rashid et al. 2026). Reducing *cofilin* (*D.m. twinstar* (*tsr*)) dosage alters both the cytoskeletal and transcriptional responses to heat, reduces the magnitude of many heat-associated transcriptional changes, and partially rescues embryo survival (Figard et al. 2019; Biel et al. 2026). These observations challenge the simple interpretation that mounting a large stress-response signature necessarily represents effective protection. Instead, strong activation of stress pathways may reflect a greater underlying molecular burden, whereas successful tolerance may depend on limiting that disruption sufficiently to maintain developmental progression. Whether distinct genetic backgrounds differ in how they balance these competing demands, and whether those differences change between acute and chronic heat exposure, remains unresolved.

Here, we compared cellularizing nuclear cycle 14 (Bownes stage 5) embryos (Bownes 1975) from an inbred Oregon-R background and the same Oregon-R-derived background outcrossed using w^1118^ under control (18°C), acute 32°C, and chronic 32°C conditions. The two backgrounds showed modest transcriptomic differences at 18°C but diverged markedly under heat. Inbred embryos exhibited a greater stress-response burden, including stronger proteostasis, DNA repair, mitochondrial, metabolic, and RNA-processing programs, yet showed poor larval hatching and adult eclosion. In contrast, genes expressed more highly in outcrossed embryos were enriched for developmental, signaling, adhesion, transport, sensory, and morphogenetic functions, and outcrossed embryos remained substantially more viable. Acute heat exposure produced the broadest background-dependent divergence, whereas chronic heat exposure retained and remodeled a smaller subset of these responses. Together, these findings associate developmental heat tolerance with the preservation of developmental gene expression programs rather than with the magnitude of canonical stress-pathway activation.

## Results

### Inbred and outcrossed embryos show divergent developmental survival after heat stress

To determine whether genetic background influences the developmental consequences of embryonic heat exposure, we measured larval hatching and adult eclosion after control (18°C), acute 32°C, and chronic 32°C heat treatments (Fig. 1A). At 18°C, both backgrounds remained highly viable and did not differ significantly at either endpoint. Larval hatching was 95.3% in the inbred background and 99.7% in the outcrossed background (P = 0.052), while adult eclosion was 90.8% and 95.3%, respectively (P = 0.158; Fig. 1B, C). Thus, under permissive conditions, the two backgrounds completed development with comparable efficiency.

**Fig. 1.**
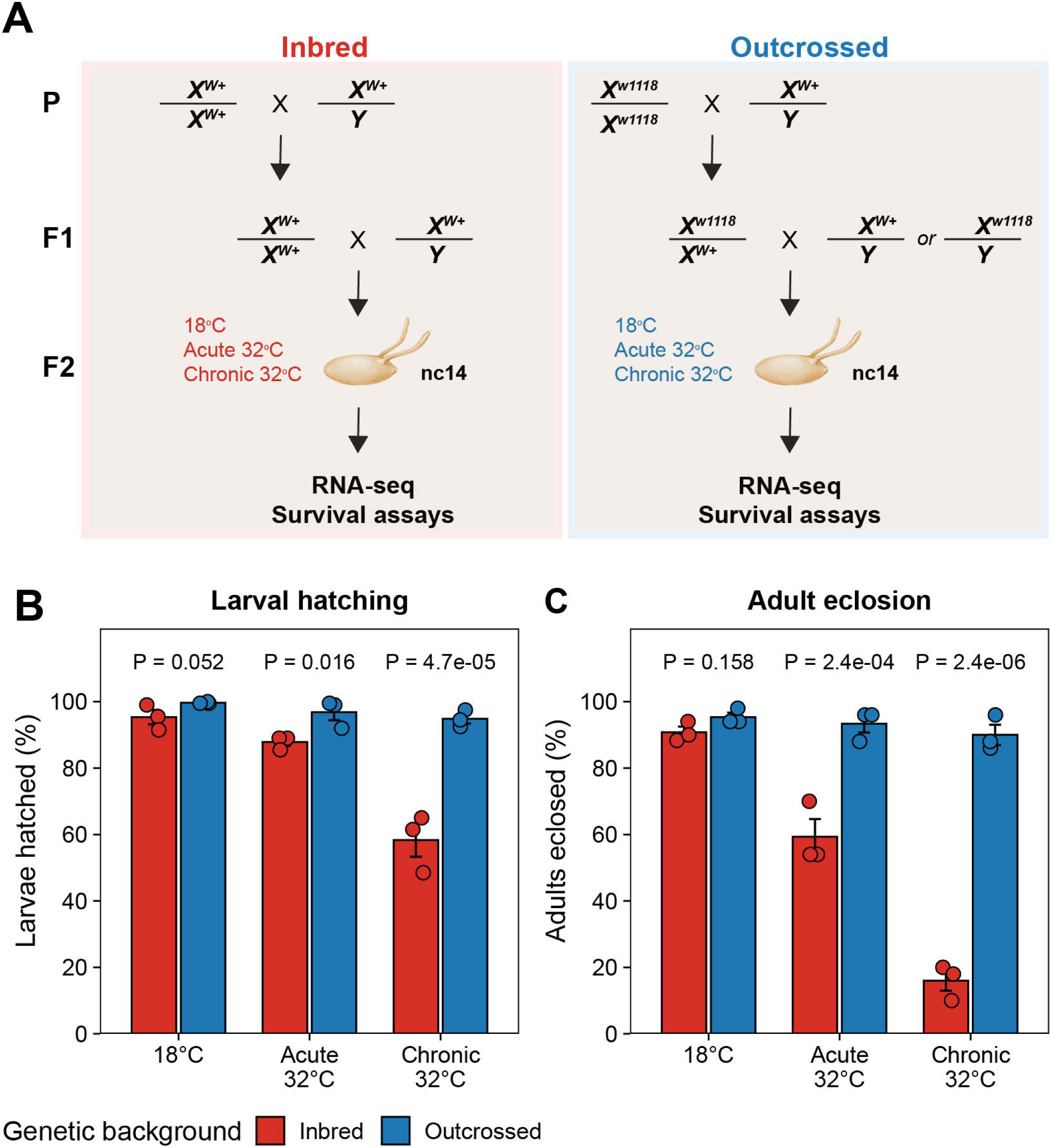
Genetic background determines developmental survival after embryonic heat stress. **A.** Experimental design. Embryos from inbred and outcrossed backgrounds were analyzed under control, acute 32°C, or chronic 32°C heat conditions. For acute heat, embryos were generated from flies maintained at 18°C and shifted to 32°C for 1.5 h before cellularization. For chronic heat, flies were maintained at 32°C for ≥ 12 h before egg collection, and embryos were subsequently laid and developed continuously at 32°C until cellularization. The embryo illustration was created in BioRender: Sokac, A. (2026) https://BioRender.com/zf6odkh. **B-C.** Larval hatching (**B**) and adult eclosion (**C**) for each genetic background and temperature condition. Both measures are expressed as a percentage of embryos scored, in independent cohorts (200 embryos per experiment for larval hatching; 50–60 for adult eclosion). Bars show the mean of three independent experiments, error bars ± SEM, and points the individual experiments. P values compare genetic backgrounds within each temperature condition, from a quasi-binomial generalized linear model fitted to replicate-level counts with experiment as a blocking factor, Holm-corrected across the three comparisons within each assay.

Heat exposure sharply separated their developmental outcomes. Following acute heat stress for 1.5 h at 32°C, 87.8% of inbred embryos hatched, compared with 96.8% of outcrossed embryos (P = 0.016). The difference was greater at the adult endpoint: 59.3% of inbred embryos reached adulthood, compared with 93.3% of outcrossed embryos (P = 2.4 × 10⁻⁴). The strongest divergence occurred under the chronic heat condition. Larval hatching declined to 58.3% in the inbred background but remained at 94.8% in the outcrossed background (P = 4.7 × 10⁻⁵). Adult eclosion fell further to 16.0% in inbred embryos, compared with 90.0% in outcrossed embryos (P = 2.4 × 10⁻⁶; Fig. 1B, C; Table S1).

Because larval hatching and adult eclosion were measured in independent cohorts, these assays do not directly quantify post-hatching mortality. Nevertheless, the background difference widened from 36.5 percentage points at hatching to 74.0 percentage points at eclosion under prolonged heat, consistent with consequences of embryonic exposure that continue to affect development after hatching. Early embryonic heat exposure has similarly been shown to influence subsequent developmental performance in *Drosophila* (Lockwood et al. 2017; Örkenby et al. 2023).

Together, these results establish a clear stress-dependent phenotypic distinction between the backgrounds. Their survival was comparable under permissive conditions, arguing against a strong constitutive developmental defect. Under heat stress, however, the outcrossed background retained high larval and adult survival, whereas the inbred background became progressively compromised across developmental endpoints. Genetic background therefore strongly influences whether embryos exposed to heat successfully complete development.

### Heat stress exposes genetic-background-dependent transcriptional states

We next asked whether the contrasting developmental outcomes were accompanied by distinct transcriptional responses. RNA sequencing was performed on nuclear cycle 14 (nc14) embryos from both backgrounds at 18°C and following acute or chronic exposure to 32°C (Fig. 2A; Table S2). Because the samples were generated across two sequencing batches, differential-expression analyses incorporated batch as a covariate. Multidimensional scaling (MDS) showed that both heat condition and genetic background contributed to transcriptomic variation, while the corresponding inbred batch controls (biological replicates) enabled the distinction of batch effects from the biological comparisons of interest (Fig. 2B).

**Fig. 2.**
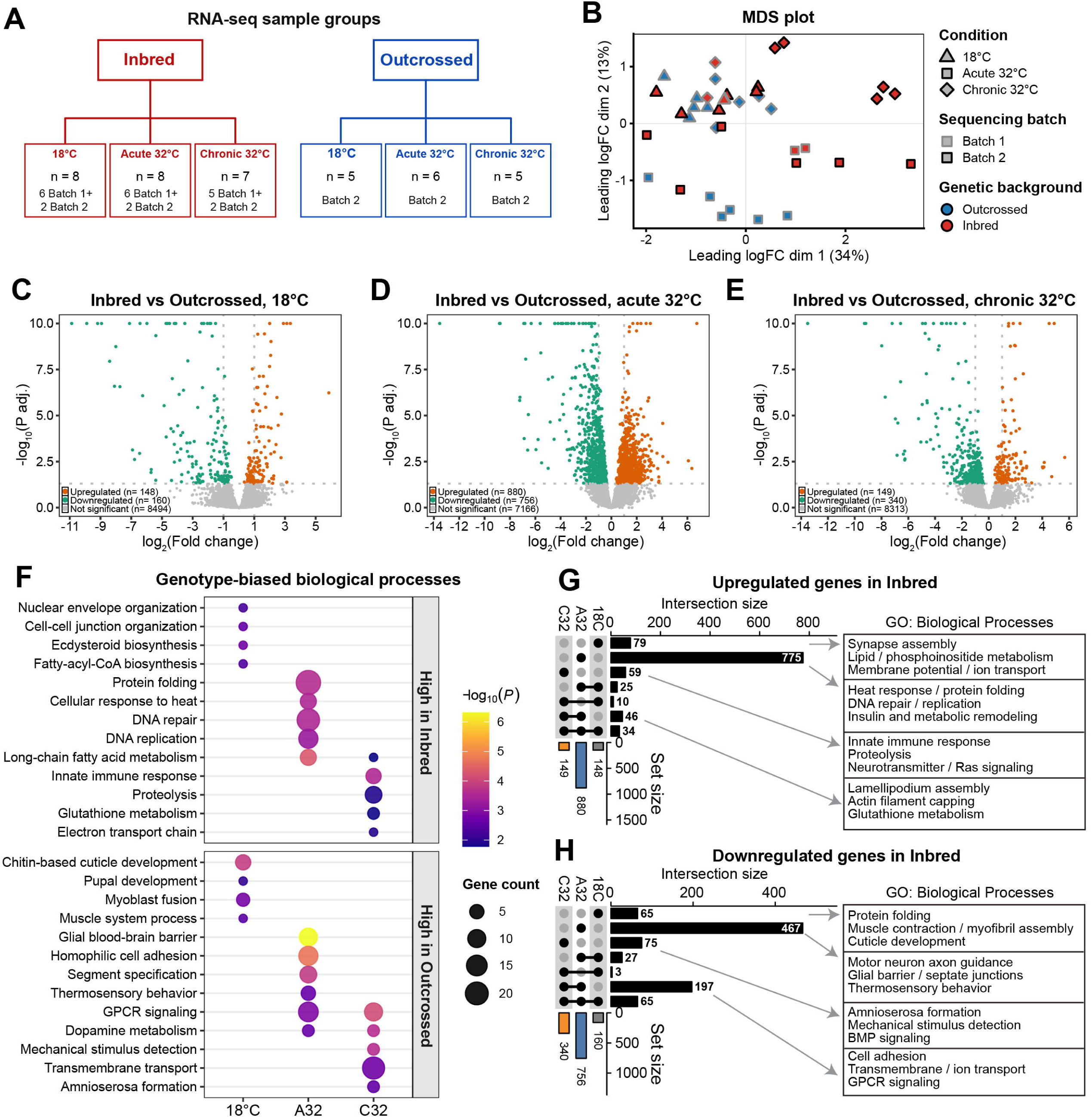
Heat stress exposes distinct gene expression programs in inbred and outcrossed embryos. **A.** RNA-seq sample groups used for batch-adjusted differential expression analysis. Stage 5 embryos were analyzed from Inbred and Outcrossed backgrounds at 18°C, acute 32°C, and chronic 32°C. **B.** MDS plot of TMM-normalized RNA-seq samples. Points are colored by genetic background, shaped by temperature condition, and outlined by sequencing batch. **C–E.** Volcano plots showing batch-adjusted differential gene expression between Inbred and Outcrossed embryos at 18°C (**C**), acute 32°C (**D**), and chronic 32°C (**E**). Positive log₂ fold change indicates higher expression in Inbred embryos; negative log₂ fold change indicates higher expression in Outcrossed embryos. **F.** Selected nonredundant GO Biological Process terms enriched among genotype-biased genes across temperature conditions. Dot size indicates gene count, and color indicates −log₁₀(P). Complete results, including BH-adjusted *P*values, are provided in Table S7. **G.** UpSet plot showing overlap of genes upregulated in Inbred embryos across 18°C, acute 32°C, and chronic 32°C. Right boxes list selected enriched GO Biological Process terms for condition-specific and heat-shared intersections. **H.** UpSet plot showing overlap of genes downregulated in Inbred embryos, corresponding to Outcrossed-high genes, across 18°C, acute 32°C, and chronic 32°C. Right boxes list selected enriched GO Biological Process terms for condition-specific and heat-shared intersections.

Three features of the design were evaluated before interpreting these comparisons. First, RNA-seq-based genotyping (Piskol et al. 2013; Danecek et al. 2021) showed substantially greater heterozygosity in the outcrossed libraries than in the inbred libraries: at a minimum depth of 20, 63.3% of autosomal exonic variant sites were called heterozygous in outcrossed libraries compared with 1.5% in inbred libraries, with no overlap between groups. Allele balance at heterozygous calls was centered near 0.5. Re-genotyping a common set of positions across the outcrossed libraries further showed extensive sharing: among 23,447 positions heterozygous in at least one library, 15,758 (67.2%) were heterozygous in all 16 libraries, and heterozygous calls were distributed across the major autosomal arms (Fig. S1A–E; Table S3). Second, because all outcrossed libraries were sequenced in the second batch, every genotype contrast was repeated using batch 2 libraries alone, where no batch correction is applied; estimated effects were highly concordant with the full model (Pearson r = 0.91–0.99), and of 4,424 significant gene-by-contrast calls only 11 differed in sign, none of which reached significance in the batch 2-only analysis (Fig. S2; Table S4, File S1). Third, because Bownes stage 5 encompasses cellularization within nuclear cycle 14, a developmental interval over which markers such as *nullo* and *bcd* change substantially (Rose and Wieschaus 1992; Surdej and Jacobs-Lorena 1998; Little et al. 2011), we tested whether the two backgrounds differed systematically in developmental position. Using both a marker-based staging index and whole-transcriptome projection onto a staged nc14 reference, we detected no such difference (P ≥ 0.088 in all comparisons; Fig. S3; Table S5) (Lott et al. 2011). The observed expression differences are therefore unlikely to be explained primarily by systematic differences in developmental staging or sequencing batch.

At 18°C, the two backgrounds differed at only 308 genes, with 148 expressed more highly in inbred embryos and 160 expressed more highly in outcrossed embryos at a false-discovery rate below 0.05 (Fig. 2C; Table S6). Heat stress substantially increased this divergence. During acute stress, 880 genes were more highly expressed in inbred embryos and 756 were more highly expressed in outcrossed embryos, producing more than five times as many background-associated differences as were detected at 18°C (Fig. 2D). The direct background comparison during chronic stress identified 149 inbred-high and 340 outcrossed-high genes (Fig. 2E). Thus, the largest separation between the backgrounds occurred during acute heat, whereas chronic heat produced fewer differences overall and was skewed toward genes expressed more highly in outcrossed embryos.

The biological composition of these gene sets changed markedly with temperature. Genes expressed more highly in inbred embryos under heat stress were enriched for protein folding, proteolysis, DNA replication and repair, glutathione metabolism, mitochondrial electron transport, long-chain fatty-acid metabolism, and other stress-associated functions (Fig. 2F; Table S7). In contrast, genes expressed more highly in outcrossed embryos were enriched for developmental and tissue-level programs, including segment specification, cell adhesion, junction and epithelial barrier organization, transmembrane transport, receptor signaling, sensory and neural functions, and morphogenesis (Fig. 2F). At 18°C, the enriched terms were more limited and included cuticle, muscle, and developmental categories, consistent with the comparatively modest baseline divergence.

Overlap analysis showed that most genotype-biased expression was condition-specific, with the largest unique component emerging during acute heat stress (Fig. 2G, H). Among genes expressed more highly in Inbred embryos, the acute-specific set was enriched for heat response and protein folding, DNA replication and repair, and insulin and metabolic remodeling, whereas the chronic-specific set was associated with innate immune signaling, proteolysis, and neurotransmitter and Ras signaling (Fig. 2G). A smaller group that remained Inbred-high during both acute and chronic heat was enriched for lamellipodium assembly, actin filament capping, and glutathione metabolism. Conversely, acute-specific Outcrossed-high genes were enriched for motor-neuron axon guidance, glial barrier and septate-junction organization, and thermosensory behavior, while chronic-specific genes were associated with amnioserosa formation, BMP signaling, and detection of mechanical stimuli (Fig. 2H). Genes that remained Outcrossed-high during both heat conditions were enriched for cell adhesion, transmembrane and ion transport, and GPCR signaling. The limited overlap with the corresponding 18°C-biased gene sets indicates that heat primarily generates new, condition-dependent differences rather than simply amplifying the transcriptional differences already present at baseline. This pattern is consistent with previous evidence that temperature can expose regulatory variation that remains comparatively buffered under permissive conditions (Chen et al. 2015).

### Acute heat elicits distinct stress-defense and developmental responses

Because the direct static comparisons indicated that background divergence was greatest during acute heat, we next examined the response to acute exposure within each background. Both backgrounds underwent extensive transcriptional remodeling. In inbred embryos, acute heat altered 1,943 genes, including 1,261 induced and 682 repressed genes. In outcrossed embryos, 2,931 genes responded, including 1,497 induced and 1,434 repressed genes (Fig. 3A, B). The outcrossed background therefore did not exhibit a globally weaker differential gene expression response. Rather, the two backgrounds differed in which genes changed and in the biological programs represented among those changes. UpSet analysis identified shared components of the acute response, together with substantial sets of genes that responded only in one background or responded in opposite directions (Fig. 3C).

**Fig. 3.**
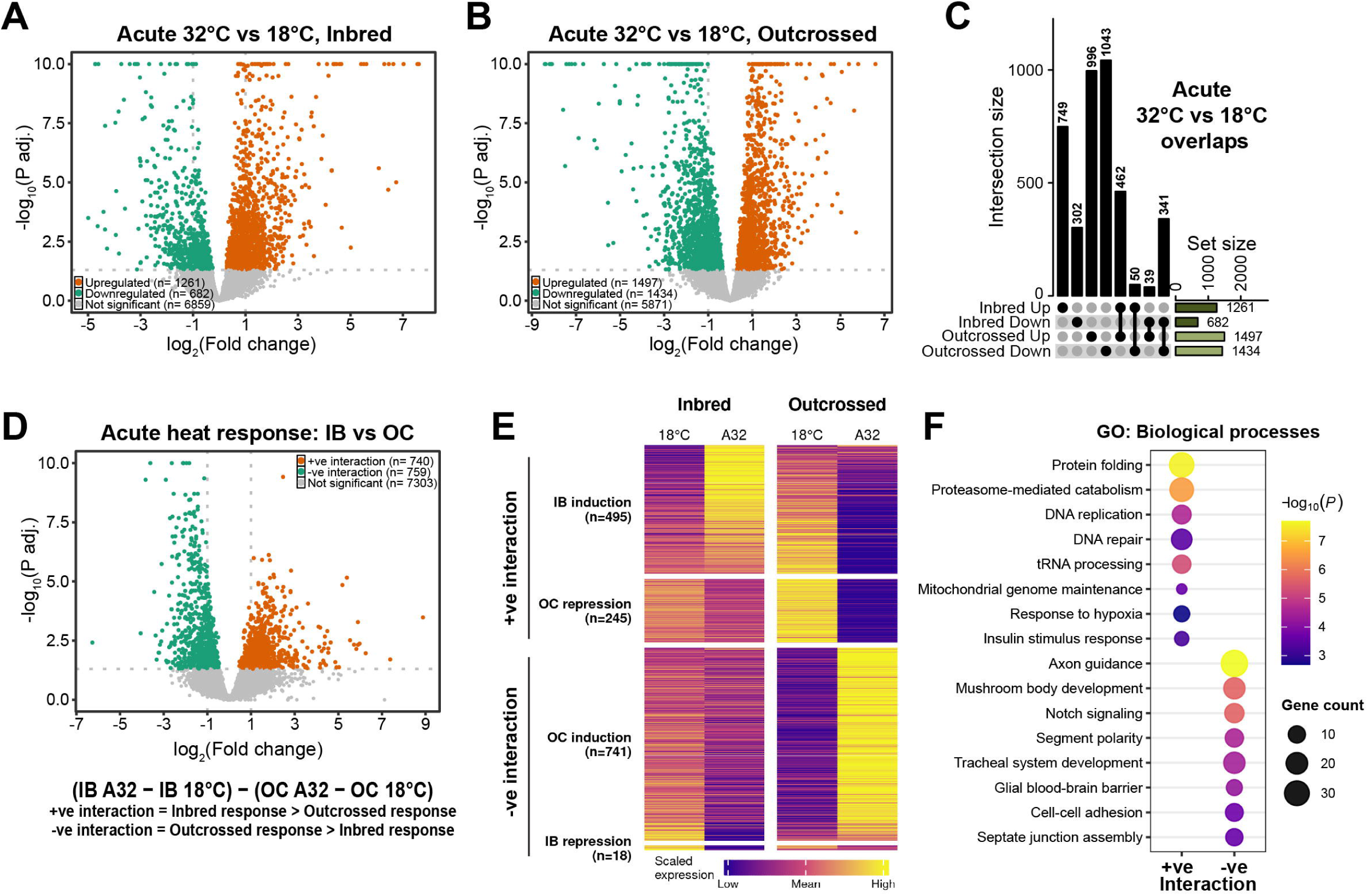
Genetic background reshapes the acute heat transcriptional response. **A-B.** Volcano plots showing acute heat-induced differential gene expression in Inbred (**A**) and Outcrossed (**B**) embryos, comparing acute 32°C with 18°C. Upregulated genes are induced by acute heat; downregulated genes are reduced by acute heat. **C**. UpSet plot showing overlap among acute heat-responsive genes in Inbred and Outcrossed embryos, separated by upregulated and downregulated genes. **D**. Acute interaction volcano plot testing genotype-dependent heat response using: (Inbred A32 − Inbred 18°C) − (Outcrossed A32 − Outcrossed 18°C). Positive interaction indicates a stronger acute response in Inbred embryos; negative interaction indicates a stronger acute response in Outcrossed embryos. **E**. Heatmap of all significant acute interaction genes using scaled group-wise mean expression. Genes are grouped by directional response class: Inbred-biased induction, Outcrossed-biased repression, Outcrossed-biased induction, and Inbred-biased repression. **F**. Selected nonredundant GO Biological Process terms enriched among positive and negative acute interaction genes.

To formally identify genes whose response to acute heat differed by background, we tested the interaction: (Inbred A32 − Inbred 18°C) − (Outcrossed A32 − Outcrossed 18°C). This analysis identified 740 positive and 759 negative interaction genes (Fig. 3D). A positive interaction indicated a greater heat-associated change in the inbred background, whereas a negative interaction indicated a greater change in the outcrossed background. The nearly equal number of positive and negative events indicate that the acute response was not shifted globally in one direction. Instead, acute heat separated the backgrounds into distinct functional response programs.

Investigation of group-wise expression patterns clarified the changes underlying these interaction effects. Among positive interaction genes, 495 were induced more strongly in inbred embryos, while 245 reflected greater repression in outcrossed embryos. Among negative interaction genes, 741 were induced more strongly in outcrossed embryos, whereas only 18 reflected greater repression in inbred embryos (Fig. 3E). Thus, much of the acute divergence arose through selective induction of different gene programs rather than through broadly opposing transcriptional shutdown.

Positive interaction genes were enriched for protein folding, proteasome-mediated protein catabolism, DNA replication and repair, mitochondrial genome maintenance, tRNA processing, hypoxia response, and metabolic signaling (Fig. 3F). These categories indicate a stronger activation of canonical stress-defense, proteostasis, repair, and metabolic-remodeling programs in inbred embryos. By contrast, negative interaction genes were enriched for axon guidance, mushroom body development, Notch signaling, segment polarity, tracheal development, cell–cell adhesion, septate junction assembly, and glial barrier organization (Fig. 3F). The outcrossed response therefore contained a stronger developmental, signaling, neural, and tissue-organization component.

Importantly, the poorer survival of inbred embryos was not associated with an inability to activate stress-response genes. Instead, these embryos carried the stronger canonical stress and damage-control signature. This distinction is consistent with previous work in the same embryonic heat-stress system, in which reducing Cofilin dosage altered the composition and magnitude of the heat-induced transcriptional response while partially improving embryo survival (Figard et al. 2019; Biel et al. 2026). Although the present data do not establish causality, they support the interpretation that a large stress-response signature can reflect a greater underlying stress burden rather than more successful adaptation.

### Chronic heat retains and remodels a selective subset of background-dependent responses

We next asked whether chronic heat exposure maintained the acute divergence or produced a distinct transcriptional state. Chronic heat caused extensive remodeling within both backgrounds, but the magnitude was greater in inbred embryos. Relative to 18°C, 2,052 genes were induced and 1,949 repressed in the inbred background, compared with 1,033 induced and 847 repressed genes in the outcrossed background (Fig. 4A, B). Overlap analysis revealed shared chronic responses but also large background-specific components (Fig. 4C). Despite the broad within-background changes, the formal chronic interaction identified only 492 genes: 160 with positive and 332 with negative interaction effects (Fig. 4D). This was substantially smaller than the 1,499 interaction genes detected after acute heat. Thus, among genes with a significant chronic interaction, stronger heat-associated changes were more often observed in the outcrossed background.

**Fig. 4.**
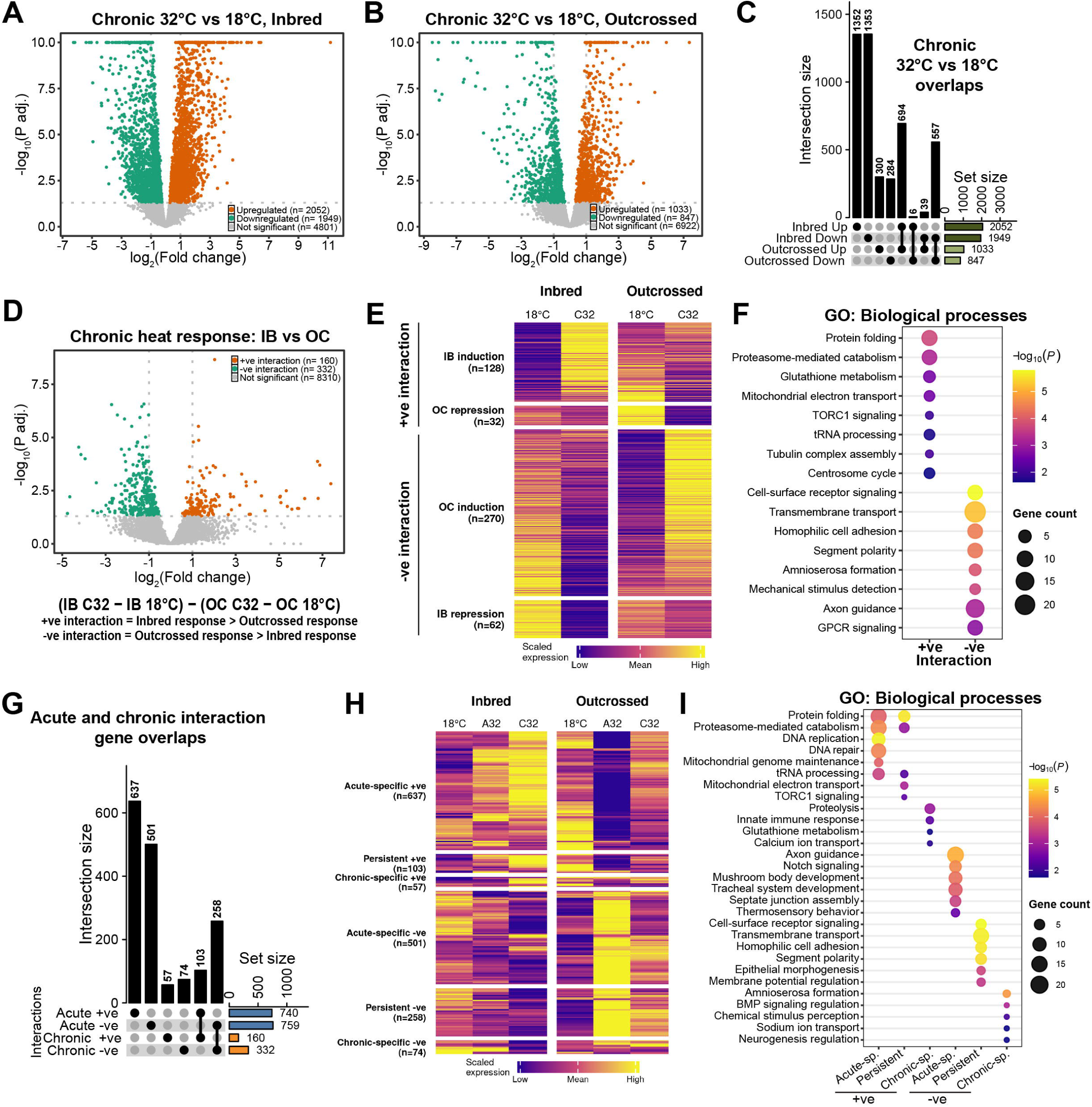
Chronic heat selectively preserves and remodels genotype-dependent transcriptional programs. **A-B.** Volcano plots showing chronic heat-induced differential gene expression in Inbred (**A**) and Outcrossed (**B**) embryos, comparing chronic 32°C with 18°C. C. UpSet plot showing overlap among chronic heat-responsive genes in Inbred and Outcrossed embryos. **D.** Chronic interaction volcano plot testing genotype-dependent chronic heat response using: (Inbred C32 − Inbred 18°C) − (Outcrossed C32 − Outcrossed 18°C). Positive interaction indicates a stronger chronic heat response in Inbred embryos; negative interaction indicates a stronger chronic heat response in Outcrossed embryos. **E.** Heatmap of significant chronic interaction genes using scaled group-wise mean expression. Genes are grouped by directional chronic-response class. Inbred-biased induction and Outcrossed-biased repression contribute to positive interaction, whereas Outcrossed-biased induction and Inbred-biased repression contribute to negative interaction. **F.** Selected nonredundant GO Biological Process terms enriched among positive and negative chronic interaction genes. G. UpSet plot comparing acute and chronic genotype-by-heat interaction genes. Acute positive, acute negative, chronic positive, and chronic negative interaction genes were compared to identify acute-specific, persistent, and chronic-specific genotype-dependent responses. For this analysis, “acute-specific,” “persistent,” and “chronic-specific” denote interaction genes reaching FDR < 0.05 under acute heat only, under both heat paradigms, or under chronic heat only, respectively. **H.** Six-group trajectory heatmap of genes with significant acute and/or chronic genotype-by-heat interaction. Columns show scaled group-wise mean expression across Outcrossed and Inbred embryos at 18°C, acute 32°C, and chronic 32°C. Genes are grouped by whether the interaction is acute-specific, persistent, or chronic-specific, and by interaction direction. I. Selected GO Biological Process terms enriched among trajectory-defined interaction gene classes from panel H. Positive trajectory classes correspond to Inbred-biased heat responses, whereas negative trajectory classes correspond to Outcrossed-biased heat responses.

Directional classification showed that positive chronic interactions consisted of 128 genes induced more strongly in inbred embryos and 32 genes repressed more strongly in outcrossed embryos. Negative interactions included 270 genes induced more strongly in outcrossed embryos and 62 genes repressed more strongly in inbred embryos (Fig. 4E). Positive interaction genes remained enriched for protein folding, proteasome-mediated catabolism, glutathione metabolism, mitochondrial electron transport, TORC1 signaling, tRNA processing, and cytoskeletal assembly (Fig. 4F). Negative interaction genes were enriched for cell-surface receptor signaling, transmembrane transport, homophilic adhesion, segment polarity, amnioserosa formation, mechanical-stimulus detection, axon guidance, and G protein-coupled receptor signaling (Fig. 4F). Chronic heat therefore preserved the broad functional distinction observed after acute exposure: stronger molecular stress and remodeling programs in inbred embryos, and stronger developmental, signaling, sensory, and transport programs in outcrossed embryos.

Comparison of the acute and chronic interaction sets revealed substantial differences in their extent and overlap. Most acute interaction genes were acute specific, including 637 positive and 501 negative genes (Fig. 4G). Nevertheless, 103 positive and 258 negative interaction genes persisted across both stress conditions. Only 57 positive and 74 negative interaction genes were unique to chronic heat. Consequently, approximately 73% of all chronic interaction genes were also significant in the acute interaction analysis. Persistent genes maintained the same interaction direction, with little evidence of switching between inbred- and outcrossed-biased responses (Fig. 4G). A six-group expression-pattern heatmap further separated acute-specific, persistent, and chronic-specific programs across both backgrounds and all three conditions (Fig. 4H). Persistent inbred-biased genes were dominated by proteostasis, mitochondrial, RNA-processing, and metabolic functions, whereas persistent outcrossed-biased genes were enriched for signaling, adhesion, transport, and developmental processes (Fig. 4I). Chronic-specific inbred-biased genes added proteolytic, redox, immune, and ion-transport functions, while chronic-specific outcrossed-biased genes included amnioserosa development, BMP signaling, sensory perception, ion transport, and neurodevelopmental regulation (Fig. 4I). Therefore, chronic heat was associated with a substantially smaller genotype-dependent program than acute heat, with extensive overlap in directionally concordant genes together with a more limited set detected only under the chronic paradigm.

Embryos undergoing cellularization contain both maternally deposited transcripts and messages transcribed from the zygotic genome, and only the latter reflect the embryo’s own transcriptional activity. We therefore classified interaction genes by transcript origin (Lott et al. 2011). Because zygotic transcripts are expressed at substantially lower levels at this stage (median 0.27 *versus* 5.78 log₂ CPM for maternal transcripts), all enrichment tests were adjusted for mean expression. Inbred-biased interaction genes were strongly enriched for maternal-zygotic transcripts –messages both deposited in the egg and transcribed by the embryo – under acute (OR = 2.98, FDR = 1.5 × 10⁻³¹) and chronic heat (OR = 1.72, FDR = 0.018) and depleted of purely zygotic transcripts (OR = 0.18 and 0.12). The heightened stress-response signature of inbred embryos therefore preferentially involved maternal-zygotic transcripts; the few purely zygotic contributors were members of the *Hsp70B* cluster (Fig. S4A, B; Table S8).

The outcrossed-biased response changed in composition with exposure duration. Purely zygotic transcripts were not enriched among outcrossed-biased acute interaction genes (OR = 0.95, FDR = 0.78) but were enriched 2.4-fold among chronic ones (OR = 2.37, FDR = 9.0 × 10⁻⁵). The same shift was evident among the enriched Gene Ontology terms: zygotic transcripts accounted for 2.1% of classified genes in outcrossed-biased acute terms but 26.3% in chronic terms, with amnioserosa formation comprising exclusively zygotic BMP pathway components (*tld*, *tsg*, *scw*, *srw*) (Fig. S4C, D). Thus, the outcrossed-biased acute interaction was dominated by transcripts classified as maternally contributed, whereas the chronic interaction included a substantially larger purely zygotic component, including genes associated with BMP-dependent dorsoventral patterning (Fig. S4C, D). Because abundance of maternally contributed transcripts at nc14 can reflect maternal deposition, transcript stability, and embryonic clearance, and chronic heat is known to disrupt maternal-transcript clearance during the maternal-to-zygotic transition (Rashid et al. 2026), the present data do not distinguish among these mechanisms.

Together, the phenotypic and transcriptomic data support a coherent model. At 18°C, the two backgrounds develop successfully and differ at relatively few genes. Heat stress reveals a much larger separation. Inbred embryos exhibit a higher transcriptional burden of proteostasis, repair, mitochondrial, redox, and metabolic programs but show progressively poorer larval hatching and adult eclosion. Outcrossed embryos retain high developmental viability and show stronger maintenance or activation of signaling, adhesion, neural, transport, and morphogenetic programs. Genetic differences in embryonic heat tolerance and stress-response tuning have also been observed among natural *Drosophila* populations (Mikucki et al. 2024). Our findings extend this principle by suggesting that successful development under heat is associated not simply with the magnitude of canonical stress-pathway activation, but with the ability to preserve developmental gene-expression programs during environmental challenge.

## Discussion

Our findings show that genetic background strongly shapes how early *Drosophila* embryos respond to heat stress. Under permissive conditions, the inbred and outcrossed backgrounds developed efficiently and differed at relatively few genes. Heat exposure, however, sharply separated both their transcriptional states and developmental outcomes. Inbred embryos exhibited broader activation of proteostasis, DNA-repair, mitochondrial, redox, metabolic, and RNA-processing programs, yet were less likely to hatch and reach adulthood. Outcrossed embryos retained higher abundance of transcripts associated with signaling, adhesion, transport, sensory function, tissue organization, and morphogenesis and showed substantially greater developmental survival (Figs. 1–4). Together, these observations suggest that genetic background shapes the balance between molecular stress-response activation and preservation of the gene-expression programs required for ongoing development.

The stronger stress-response signature in inbred embryos should not be interpreted as evidence of a more effective protective response. Inbred embryos were clearly capable of activating canonical stress-associated programs, but broader activation did not translate into greater developmental survival. Such broad activation may instead reflect, at least in part, the degree of molecular perturbation experienced by the embryo. A similar disconnect has been observed previously: inbred flies showed elevated Hsp70 expression despite reduced heat resistance (Pedersen et al. 2005). Individual protective factors can nevertheless improve thermal tolerance; increased maternal loading of Hsp23, for example, enhances embryo survival after acute heat exposure and reduces subsequent defects in larval performance (Lockwood et al. 2017). The relevant distinction may therefore be between protective responses that successfully limit damage and a broad emergency program associated with substantial proteotoxic, metabolic, or structural disruption. Under this model, the expanded stress-defense signature in inbred embryos reflects a higher stress burden, whereas the more restrained signature in outcrossed embryos reflects reduced disruption rather than a failure to recognize heat.

Preservation of development-associated transcripts provides a complementary measure of this difference. Cellularizing embryos are establishing body-patterning information while preparing for the morphogenetic movements of gastrulation, processes that require coordinated transcription, signaling, adhesion, cytoskeletal organization, and tissue specification (Stathopoulos and Newcomb 2020; Sokac et al. 2023). Although *Drosophila* embryogenesis can adjust its rate across a broad temperature range while preserving the relative timing of developmental events (Kuntz and Eisen 2014), severe thermal perturbation can exceed this buffering capacity. In our study, greater abundance of developmental, neural, epithelial, adhesion, transport, and morphogenetic transcripts in outcrossed embryos was associated with higher larval hatching and adult eclosion. Conversely, the larger stress-response burden in inbred embryos accompanied substantial developmental attrition after chronic heat exposure: approximately 58% hatched, but fewer than 20% of the original cohort reached adulthood (Fig. 1). Early embryonic heat exposure has previously been shown to affect later developmental performance (Graziosi et al. 1983; Walter et al. 1990; Wang and Lindquist 1998; Lockwood et al. 2017; Figard et al. 2019; Gibbs et al. 2025; Biel et al. 2026), consistent with the idea that disruption during this restricted developmental window can have consequences after the initial exposure has ended.

Transcript-origin analysis further refines the interpretation of these developmental signatures. Under acute heat, outcrossed-biased interaction genes were predominantly maternally derived, with no enrichment for purely zygotic transcripts (Fig. S4). At the Bownes stage 5, the abundance of these transcripts reflects maternal provisioning, transcript stability, and embryonic clearance during the maternal-to-zygotic transition (Bashirullah et al. 1999; Tadros and Lipshitz 2009). Higher abundance in outcrossed embryos could therefore arise through any of these processes, which cannot be distinguished in the present design. Under chronic heat, the outcrossed-biased response contained a larger zygotic component, including patterning genes associated with BMP-dependent dorsoventral development. This finding suggests that background-dependent differences extend to the embryo’s own transcriptional programs under sustained exposure. However, because the prolonged-heat condition also included maternal exposure before and during egg laying, maternal and embryonic contributions remain partly confounded. Reciprocal crosses and allele-specific expression analysis would be required to separate these effects.

The two heat paradigms also revealed different extents of background-dependent transcriptional divergence. Acute heat produced the largest set of genotype-by-heat interactions, whereas fewer interaction genes were detected after chronic exposure (Figs. 3 and 4). Genes significant under both conditions generally retained the same interaction direction, indicating that part of the background-dependent response was shared across treatments. Other genes reached significance only under acute or chronic heat and were associated with distinct stress, metabolic, transport, signaling, and developmental processes. These categories should be interpreted descriptively, because significance under one condition but not the other does not establish that the response differs statistically between treatments. A direct contrast between the acute and chronic genotype-by-heat interaction effects would be required to formally define treatment-specific responses. Moreover, because the two paradigms differ in maternal thermal history as well as embryonic exposure, their transcriptional differences cannot be attributed solely to exposure duration.

The exposure of genetic-background differences by heat is consistent with broader evidence for genotype–environment interactions in Drosophila. Oregon-R and Samarkand show relatively little expression divergence at 18°C but substantially greater divergence at temperature extremes, supporting the idea that environmental stress can reveal regulatory variation that remains buffered under permissive conditions (Chen et al. 2015). Inbreeding and elevated temperature have also been shown to interact in their effects on metabolic and heat-responsive gene expression, with larger expression changes accompanying reduced fitness in inbred lines (Kristensen et al. 2006). More recently, embryos from tropical and temperate populations were shown to differ in heat tolerance and oxidative-stress pathway regulation (Mikucki et al. 2024). Our study extends these observations by showing that heat-exposed backgrounds differ not only in the extent of their transcriptional response, but also in whether that response is dominated by molecular stress programs or by the continued abundance of development-associated transcripts.

The amplification of background-dependent differences under heat is broadly consistent with genetic variation that is comparatively buffered under permissive conditions becoming consequential during stress. The outcrossed embryos in this study were generated from OreR and w^1118^ parental backgrounds but constituted segregating second-generation populations. Increased heterozygosity could contribute to the observed phenotype by masking partially recessive variants (Charlesworth and Willis 2009), but effects of individual parental alleles, dominance, epistasis and recombination cannot be distinguished in the present design. Resolving these contributions will require reciprocal crosses and multiple independently derived crosses.

Several limitations define the scope of these conclusions. The study compares one inbred OreR background with outcross-derived populations generated from OreR and w1118 and therefore does not establish a universal effect of inbreeding, outcrossing, or heterozygosity. The observed differences could reflect heterozygosity, segregation of parental alleles, dominance, epistasis, maternal genetic effects, or combinations of these factors. Because a substantial fraction of the acute background-dependent signal involves maternally derived transcripts, the relative contributions of maternal provisioning, transcript stability, and embryonic clearance also remain unresolved. Importantly, the mothers producing the two experimental backgrounds were also genetically different, so maternal-genotype effects are intrinsic to this comparison. Bulk RNA sequencing measures the average state of the embryo and cannot determine whether developmental programs are preserved uniformly or within specific tissues. Spatial analysis of developmental transcripts and imaging of epithelial, nervous-system, or morphogenetic development would directly test whether the expression differences correspond to improved tissue organization. Finally, the association between developmental-transcript preservation and survival does not demonstrate causality. The most parsimonious interpretation is that genetic background alters the immediate molecular and developmental consequences of heat, which then influence the probability of completing later stages of development.

Overall, the results support a model in which the two backgrounds occupy different positions along a stress–development axis during heat exposure. Inbred embryos exhibit a broad molecular stress response accompanied by substantial developmental failure, whereas outcrossed embryos show a lower apparent stress burden, greater preservation of development-associated transcripts, and a higher probability of completing development. Genetic background therefore influences not only the transcriptional response to heat, but also how effectively that response is integrated with the ongoing demands of embryogenesis.

## Materials and Methods

### Fly stocks and husbandry

*Drosophila melanogaster* stocks were maintained on Fly Food R (Lab-Express, Ann Arbor, MI, USA) under standard laboratory conditions. Oregon-R (OreR) was used as the inbred background. To generate the outcross-derived experimental groups, virgin w1118 females were crossed with OreR males. For RNA-sequencing experiments, virgin female F1 progeny were crossed with F1 sibling males, and F2 embryos from this cross were analyzed. For survival experiments, virgin female F1 progeny were crossed with OreR males, and embryos from this backcross were analyzed. Thus, the outcrossed experimental groups were derived from the same parental backgrounds but were segregating populations. Throughout the manuscript, “inbred” and “outcrossed” refer operationally to these experimental backgrounds.

### Embryo collection and heat-stress exposure

Embryo collection and heat-stress exposure were performed as described previously (Biel et al. 2026; Rashid et al. 2026). Embryos were collected on apple juice agar plates attached to collection cups. Before embryo collection, flies were acclimated to the starting temperature: collection cups for the control and acute-stress groups were maintained at 18°C for at least 48 h, whereas cups for the chronic-stress group were maintained at 32°C for 12-24 h. For the 18°C control condition, flies were allowed to lay eggs for 2 h, after which collection plates were incubated for an additional 3.5 h at 18°C. For acute heat stress, flies laid eggs for 2 h at 18°C, plates were incubated for an additional 20 min at 18°C, and embryos were then shifted to 32°C for 1.5 h. For chronic heat stress, flies laid eggs for 1 h at 32°C, after which plates remained at 32°C for an additional 1.5 h. Thus, in the acute paradigm embryos first encountered 32°C during embryonic development, whereas in the chronic paradigm maternal flies were exposed to 32°C before egg laying and embryos subsequently developed continuously at 32°C until collection. Embryos at Bownes stage 5 (nc14) were hand-selected under a dissecting microscope.

### Larval hatching and adult eclosion assays

Developmental survival was measured at larval hatching and adult eclosion. Embryos were hand-selected at Bownes stage 5 following acute heat-stress, or chronic heat-stress treatment and transferred to 22°C for the remainder of development, alongside non-stressed controls reared at 18°C until Bownes stage 5 and then also shifted to 22°C. Larval hatching and adult eclosion were scored on independent cohorts of hand-selected embryos, and both are expressed as a percentage of embryos scored. For larval hatching assays, stage-matched embryos with intact chorions were aligned in rows on fresh agar plates. Plates were placed in humidified boxes and maintained at 22°C for 48 h. Empty eggshells and unhatched embryos were then counted, and larval hatching was calculated as the percentage of selected embryos that successfully hatched. For adult eclosion assays, stage-matched embryos were arranged on approximately 0.8 × 0.8-cm agar blocks. Each agar block was transferred, embryo side upward, into a vial containing fresh Fly Food R. Vials were maintained at 22°C for 14 days, after which live adults and non-eclosed pupae were counted. Vials containing non-eclosed pupae were retained for an additional 3 days to include delayed eclosion. Pupae that failed to eclose by day 17 were visually examined and scored as dead. Survival experiments were performed using three independent biological replicates per genetic background and temperature condition, with 200 embryos per replicate for larval hatching and 50 embryos per replicate for adult eclosion (60 in one replicate; 150–160 embryos per group in total). Bars show the mean percentage across biological replicates, error bars the standard error of the mean, and points the individual replicates. Because survival was measured on replicate plates, statistical analysis was performed at the level of the independent biological replicate rather than on pooled embryo counts. Success and failure counts were modelled by quasi-binomial generalized linear model with a logit link (cbind(alive, dead) ∼ replicate + genotype × temperature), with replicate included as a fixed blocking factor because each biological replicate contained all six genotype– temperature groups. The quasi-binomial family estimates a dispersion parameter rather than fixing it at one, thereby accommodating variation between replicates beyond binomial sampling error. Because dispersion is estimated rather than known, inference used the t distribution on residual degrees of freedom. Genetic backgrounds were compared within each temperature condition using estimated marginal means, and the resulting three P values were corrected by the Holm procedure separately for each assay. Exact adjusted P values are reported in Figure 1; replicate-level counts and full statistical output are provided in Table S1. Analyses were performed in R (version 4.6.1) using the emmeans package (version 2.0.4).

### Embryo collection and RNA isolation for RNA sequencing

Embryos for RNA sequencing were collected using the same temperature paradigms described above. Water, collection tools, agar plates, and other reagents were pre-equilibrated to the corresponding experimental temperature whenever possible. Bownes stage 5 embryos were hand-selected under Halocarbon 27 oil, transferred to low-retention tubes, snap-frozen in liquid nitrogen, and stored at −80°C. Total RNA was isolated using the RNeasy Plus Micro Kit (Qiagen, Germantown, PA, USA) according to the manufacturer’s instructions. RNA quality was evaluated using an AATI Fragment Analyzer (Advanced Analytical Technologies, Ames, IA, USA) by measuring RNA quality number and the 28S/18S ribosomal RNA ratio. The final analysis included five outcrossed and eight inbred biological replicates at 18°C, six outcrossed and eight inbred replicates following acute heat stress, and five outcrossed and seven inbred replicates following chronic heat stress. Samples were generated in two experimental and sequencing batches. The first batch contained the original inbred samples. The second batch contained all outcrossed samples together with two additional OreR control samples for each temperature condition, which served as bridging controls for estimating batch effects.

### RNA-sequencing library preparation

Stranded messenger RNA libraries were generated by the DNA Services Laboratory at the Roy J. Carver Biotechnology Center, University of Illinois Urbana-Champaign, using the KAPA mRNA HyperPrep Kit (Roche, Indianapolis, IN, USA). Polyadenylated RNA was isolated from 200 ng of total RNA per sample and fragmented at 94°C for 4 min. First-strand complementary DNA was synthesized using random hexamers and SuperScript II Reverse Transcriptase (Invitrogen/Thermo Fisher Scientific). Double-stranded complementary DNA was end-repaired, 3′ A-tailed, and ligated to universal adapters. Libraries were amplified for 12 PCR cycles using Twist unique dual-index primers (Twist Bioscience, San Francisco, CA, USA) and diluted to a final concentration of 5 nM. Libraries were sequenced as 150-bp paired-end reads on an Illumina NovaSeq 6000 SP flow cell. Base-call files were demultiplexed and converted to adapter-trimmed FASTQ files using bcl2fastq v2.20.

### Read alignment and gene-level quantification

Sequencing-read quality was evaluated using FastQC (version 0.11.9). Paired-end reads were aligned to the *D. melanogaster* reference genome from FlyBase (FASTA: dmel-all-chromosome-r6.50.fasta, GTF: dmel-all-r6.50.gtf) (Larkin et al. 2021) using STAR (version 2.7.11b) (Dobin et al. 2013). Coordinate-sorted BAM files were generated during alignment. Sorted bam files were then indexed using SAMtools index (Li et al. 2009). Gene-level read counts were obtained using HTSeq-count (version 2.0.2) (Anders et al. 2015) in reverse-stranded mode against the corresponding FlyBase gene annotation. Meta-count rows representing unassigned, ambiguous, low-quality, nonaligned, or nonunique reads were removed before downstream analysis. FlyBase gene identifiers were mapped to gene symbols using the Bioconductor `org.Dm.eg.db` annotation package.

### Differential gene-expression analysis

Differential gene-expression analysis was performed in R using edgeR [version 3.40.2] (Robinson et al. 2010). All samples were analyzed in a single model to ensure common expression filtering, normalization, and dispersion estimation. The six genotype–condition combinations were represented as separate groups: Outcrossed 18°C, Outcrossed acute 32°C, Outcrossed chronic 32°C, Inbred 18°C, Inbred acute 32°C, and Inbred chronic 32°C. Genes with insufficient expression were removed using `filterByExpr`. Library-composition differences were normalized using the trimmed mean of M-values method (Robinson and Oshlack 2010). Multidimensional scaling was performed from normalized expression values using leading Log₂-fold-change distances. Negative-binomial generalized linear models were fitted using the design: `∼ 0 + genotype_condition_group + sequencing_batch.` The additional OreR controls included in the second sequencing batch provided the cross-batch replication needed to estimate the batch coefficient. Dispersion was estimated using the robust option in edgeR, and likelihood-ratio tests were used to evaluate prespecified contrasts. The batch coefficient was included as a nuisance covariate and was not interpreted biologically. The primary contrasts included direct comparisons between the backgrounds at each temperature and acute and chronic heat responses relative to 18°C within each background. Genotype-dependent heat responses were tested using difference-in-differences contrasts:

Acute interaction = (Inbred A32 − Inbred 18°C) − (Outcrossed A32 − Outcrossed 18°C)
Chronic interaction = (Inbred C32 − Inbred 18°C) − (Outcrossed C32 − Outcrossed 18°C)

For these contrasts, a positive Log₂-fold change indicated a stronger heat-associated response in the inbred background, whereas a negative value indicated a stronger response in the outcrossed background. P values were corrected using the Benjamini–Hochberg procedure. Genes with a false-discovery rate below 0.05 were considered differentially expressed. No minimum absolute fold-change threshold was required for significance; genes were classified as up- or downregulated according to the sign of the estimated Log₂-fold change. Heatmap values show TMM-normalized log₂CPM with batch adjustment; statistical comparisons were batch-adjusted as described above. All differential gene expression results provided in Table S6.

### RNA-seq-based characterization of heterozygosity

To characterize heterozygosity in the experimental backgrounds, variants were called from the RNA-seq alignments used for expression analysis. For the per-library analysis in Fig. S1A–C, variant calling was restricted to merged autosomal exon intervals derived from the FlyBase r6.50 annotation; sex chromosomes were excluded. Variants were called independently for each library using bcftools v1.24 (mpileup, minimum mapping quality 20, minimum base quality 20, maximum depth 500; multiallelic caller, variant sites only) (Danecek et al. 2021), and genotypes were retained at QUAL ≥30 and minimum depths of 10, 20, or 30. The heterozygous-call fraction was calculated as the proportion of called variant sites classified as heterozygous. Allele balance at heterozygous calls was evaluated as an additional quality-control measure.

For the site-sharing and spatial analyses in Fig. S1D–E, the union of variant positions detected across the 16 outcrossed libraries was compiled and every library was re-genotyped at these positions without restricting output to variant sites, thereby recovering explicit homozygous-reference calls. Analyses were restricted to the major autosomal arms 2L, 2R, 3L, and 3R. Calls were required to have depth ≥20 and a valid diploid genotype; any diploid genotype containing two different alleles, including multiallelic calls such as 1/2, was classified as heterozygous. Because multiple variant records could occur at the same genomic coordinate, any position represented by more than one record in any library was excluded from the common-site universe. Only positions with a valid call in all 16 outcrossed libraries were retained. For Fig. S1D, the number of distinct libraries called heterozygous was tabulated at each retained position.

For Fig. S1E, heterozygous-call fractions were calculated in 1-Mb windows using the same common-site universe. The displayed analysis was restricted to windows containing at least 50 common callable positions; an analysis using a minimum of 20 positions per window is provided as a sensitivity analysis in Table S3. As an internal check, heterozygous-call fractions were compared among temperature conditions within each window using Kruskal–Wallis tests with Benjamini–Hochberg correction; no window showed a significant condition association. Because the common site universe was ascertained from the union of positions initially identified as variants in the outcrossed libraries, positions homozygous-reference in all libraries were not represented. The resulting values therefore are not unbiased estimates of genome-wide heterozygosity. Within a given window, all libraries were evaluated at the same adequately covered positions, whereas differences among genomic windows are interpreted descriptively because the composition and ascertainment of evaluated sites differ among intervals.

### Batch sensitivity analysis

To test whether genotype effects depended on the batch adjustment, the six-group model was refitted using only Batch 2 libraries (design `∼0 + group`, without a batch term), retaining the gene set from the primary analysis and recomputing normalization factors and dispersion estimates on the subset. Log₂ fold changes were compared with the primary model for genes reaching FDR < 0.05 in the primary analysis. Because Batch 2 contains only two inbred replicates per condition, this analysis is underpowered for gene discovery and was used to assess concordance of effect estimates rather than to call genes. Genes with discordant effect signs were annotated and cross-referenced against the Gene Ontology enrichment results.

### Developmental staging control

Because Bownes stage 5 corresponds to cellularization within nuclear cycle 14, during which the embryonic transcriptome changes rapidly, we assessed whether inbred and outcrossed embryo collections differed in developmental position using two independent approaches. First, a marker-based staging index was computed from three panels of transcripts with known directional behaviour across nc14: early zygotic (*slam*, *nullo*, *bnk*, *halo*), late zygotic (*sna*, *twi*, *sog*, *zen*, *fog*, *T48*, *trbl*, *htl*, *ths*), and maternally deposited transcripts subject to degradation during the maternal-to-zygotic transition (*bcd*, *nos*, *osk*, *tor*, *twe*, *cort*, *exu*, *stg*). Hsp83 was excluded from the maternal panel despite being a canonical degradation target because it is heat inducible. Two additional early markers (*sry-α*, *frs*) did not pass expression filtering and were excluded. Each transcript was z-scored across samples, panel scores were computed as the mean z-score across constituent transcripts, and the staging index was defined as the late zygotic score minus the early zygotic score, such that higher values correspond to later positions within nc14. Second, samples were projected onto a staged nc14 reference derived from single-embryo RNA-seq (Table S2) (Lott et al. 2011). Reference expression was averaged across sexes and replicates within each staged bin and log₂-transformed, and gene symbols were mapped to FlyBase identifiers (99.5% mapped; duplicate identifiers were collapsed retaining the most highly expressed representative), yielding 12,080 genes of which 8,063 overlapped the expressed gene set. Analysis was restricted to bins nc14A-D. The 2,000 genes with the greatest variance across these bins were retained, and expression was z-scored independently within the reference and within the experimental data so that only the relative shape of each gene’s profile was compared. Spearman correlations were computed between each sample and each reference bin, and a continuous pseudostage was derived as the correlation-weighted mean bin index. For both measures, the additive sequencing-batch effect was removed by linear model (score ∼ group + batch) prior to comparison and inbred and outcrossed samples were compared within each temperature condition by Welch’s two-sample t-test. No systematic difference in developmental position was detected between the backgrounds.

### Maternal and zygotic transcript classification

Transcripts were classified as maternal, zygotic or maternal-zygotic using single-embryo RNA-seq assignments across nuclear cycles 10–14 (Table S2) (Lott et al. 2011). Gene symbols were mapped to FlyBase identifiers using org.Dm.eg.db, querying SYMBOL and then ALIAS. Because zygotic transcripts are expressed at lower levels than maternal transcripts at this stage, expression filtering removed a larger proportion of zygotic (38.1%) than maternal (3.7%) genes; classifications were available for 7,557 of 8,802 expressed genes (85.9%).

Enrichment of each origin class within interaction gene sets was assessed relative to the expressed background by Fisher’s exact test with Benjamini–Hochberg correction. Because median expression differed markedly among classes (5.78, 3.62 and 0.27 Log₂CPM for maternal, maternal-zygotic and zygotic transcripts respectively) and interaction contrasts have the lowest statistical power in this design, enrichment was additionally tested by logistic regression including mean log₂CPM as a covariate. Expression-adjusted odds ratios are reported throughout. Primary analyses were restricted to genes with an assigned classification, since unassigned genes were the lowest-expressed group in the dataset (median − 0.97 log₂ CPM) and treating them as negatives would import an expression bias; analyses including unassigned genes are provided in Table S8 as a sensitivity check. Composition of Gene Ontology terms was determined from the overlapping gene identifiers reported for the terms displayed in Figs. 3 and 4. For the aggregate acute-versus-chronic comparison of outcrossed-biased terms, each gene was counted once per contrast, even if it contributed to multiple displayed terms.

### Overlap and expression-pattern analyses

Gene-set intersections were calculated using unique FlyBase gene identifiers. UpSet plots were generated using the ComplexHeatmap package (Gu et al. 2016), with each gene assigned to its exact or “distinct” intersection. For interaction heatmaps and gene-level expression panels, the additive sequencing-batch effect was removed from TMM-normalized log₂ CPM using limma removeBatchEffect (Ritchie et al. 2015), protecting the six genotype–condition group means, and adjusted values were averaged across biological replicates within each group. Values were then standardized separately for each gene across the plotted group means. Batch adjustment was applied for display only; all statistical comparisons derive from the edgeR model described above. Significant interaction genes were classified according to the sign of the interaction coefficient and the direction of the within-background heat response. Acute and chronic interaction genes were further grouped as acute-specific, persistent, or chronic-specific. Persistent genes were significant in both interaction contrasts and retained the same direction during acute and chronic heat.

### Gene Ontology enrichment analysis

Gene Ontology Biological Process enrichment was performed separately for the indicated upregulated, downregulated, positive-interaction, negative-interaction, and trajectory-defined gene sets using PANGEA (Hu et al. 2023). Analyses used the *D. melanogaster* Gene Ontology Biological Process collection and the complete set of genes retained after expression filtering as the statistical background. Complete enrichment results, including nominal P values and Benjamini–Hochberg-adjusted values, are provided in Table S7. For figure presentation, closely related or redundant terms were consolidated, and a representative nonredundant subset was selected to summarize the dominant biological programs. Dot size represents the number of genes overlapping each term, and color represents −log_10_(P). All displayed terms were required to pass P < 0.05.

## Data and code availability

Inbred OreR RNA-seq dataset is available through the NCBI SRA under BioProject PRJNA1196936. Normalized expression values are provided in File S2. Analysis code will be made publicly available upon publication. The outcrossed RNA-seq libraries and additional OreR batch-control libraries are deposited in the NCBI SRA under BioProject PRJNA1368972 and will be released upon publication. Complete differential-expression tables, interaction results, Gene Ontology outputs, and source data underlying the present analyses are provided as supplementary files. Normalized expression values are provided in File S2. Analysis code for the analyses reported here will be made publicly available upon publication.

## Acknowledgments

The embryo illustration in Fig. 1A was created in BioRender (Sokac, A. (2026). BioRender.com/zf6odkh).

---

**Fig. S1.**
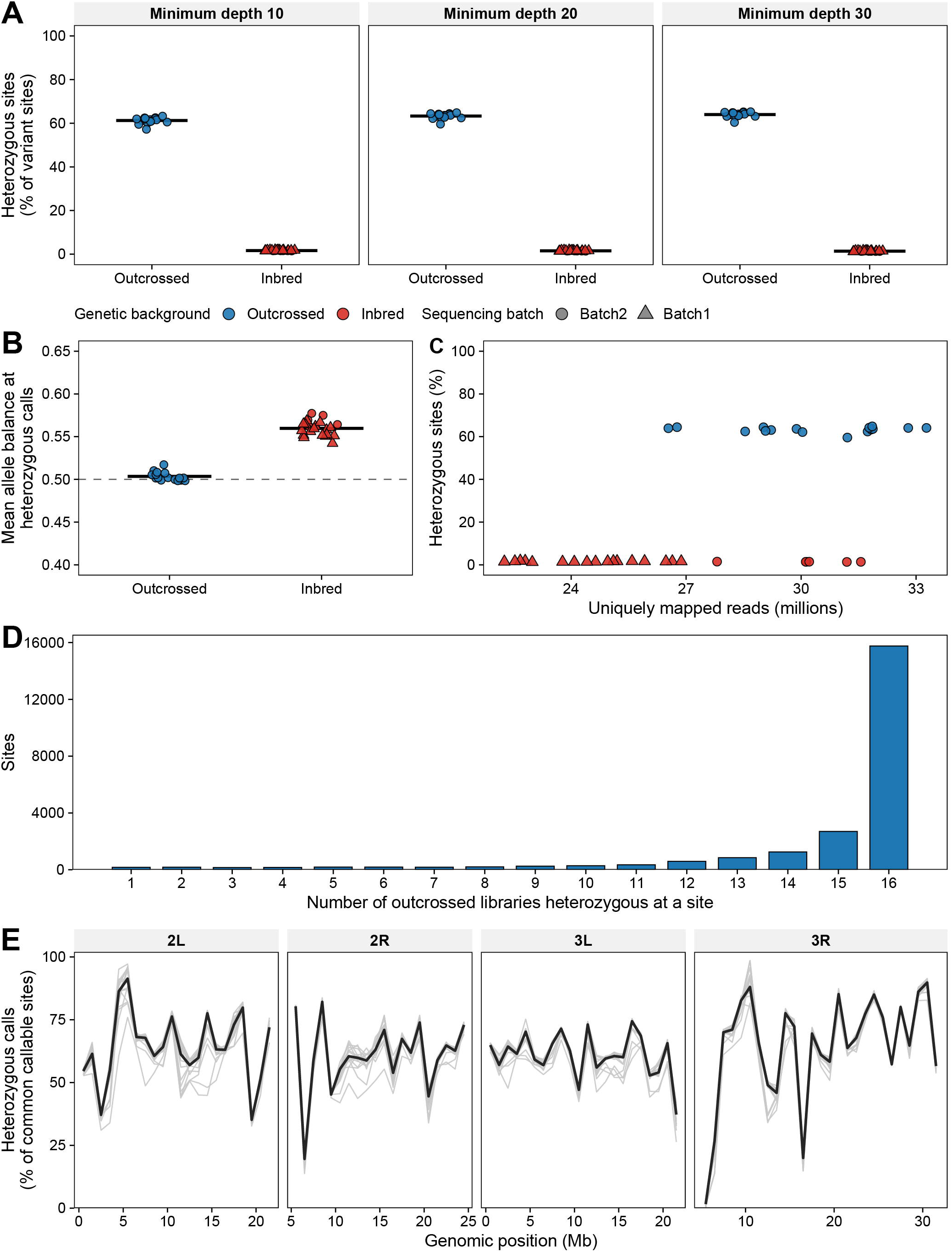
RNA-seq-based heterozygosity analysis distinguishes the inbred and outcrossed backgrounds. **A.** Proportion of autosomal exonic variant sites called heterozygous per library at three minimum-depth thresholds; points represent libraries and crossbars group means. At minimum depth 20, the mean heterozygous-call fraction was 1.5% in inbred libraries (n = 23) and 63.3% in outcrossed libraries (n = 16), with no overlap between groups. **B.** Mean allele balance at heterozygous calls. The outcrossed libraries were centered near 0.5 (dashed line), consistent with balanced representation of the two called alleles. **C.** Heterozygous-call fraction plotted against library size, showing no systematic dependence on sequencing depth. **D.** Distribution of the number of outcrossed libraries called heterozygous at common positions on the major autosomal arms with adequate coverage in all 16 libraries. Of 23,447 positions heterozygous in at least one outcrossed library, 15,758 (67.2%) were heterozygous in all 16, demonstrating extensive shared heterozygosity among the sampled libraries. **E.** Spatial distribution of RNA-seq-derived heterozygous calls across 1-Mb windows of the major autosomal arms. Only windows containing at least 50 common callable positions are shown. Light-gray traces represent individual libraries, and the dark trace represents the across-library median. Within each window, all libraries were evaluated at the same adequately covered positions. Because the evaluated positions were ascertained from the union of variants detected across the outcrossed libraries, absolute values are not unbiased estimates of genomic heterozygosity and differences among genomic windows are interpreted descriptively. The corresponding analysis using a 20-site minimum is provided in Table S3.

**Fig. S2.**
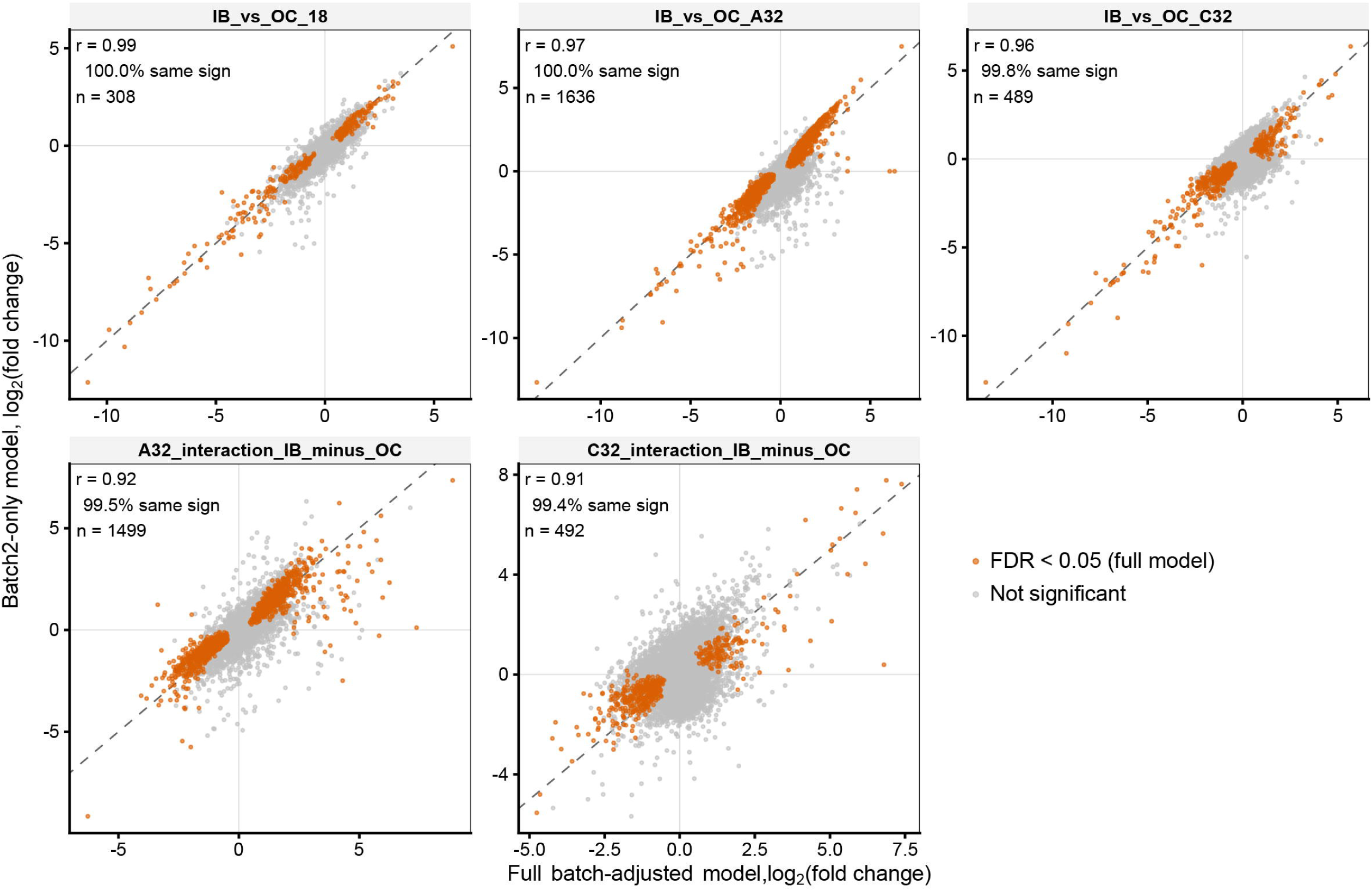
Genotype-effect estimates are concordant in the Batch 2-only analysis. Log₂ fold changes from the primary batch-adjusted model (x-axis) compared with a model fitted to batch 2 libraries only, in which no batch correction is applied (y-axis), for three genotype contrasts and two genotype-by-heat interaction contrasts. Orange, FDR < 0.05 in the primary model; grey, all other genes; dashed line, y = x. Insets give Pearson correlation, percentage sign concordance, and the number of genes significant in the primary model. Estimates were highly concordant (*r* = 0.91– 0.99); of 4,424 significant gene-by-contrast calls, 11 differed in effect sign and none reached significance in the batch 2-only model (all FDR > 0.34). Because batch 2 contains only two inbred replicates per condition, this analysis is power-limited by design and assesses agreement of effect estimates rather than significance; interaction contrasts show modest attenuation (slopes 0.74– 0.83) consistent with selection on significance in the primary model.

**Fig. S3.**
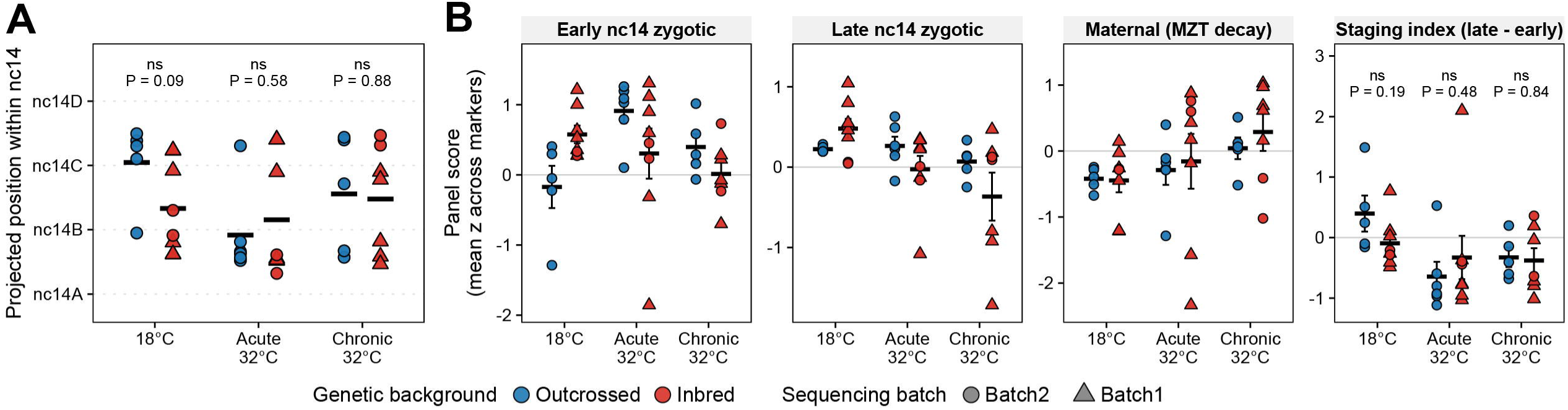
Inbred and outcrossed embryos were collected at equivalent developmental positions within nc14. **A.** Projected position within nuclear cycle 14 for each RNA-seq library, derived by correlating the 2,000 most dynamic genes against a staged nc14 reference time course (Lott *et al*. 2011) and expressed as a correlation-weighted mean bin index. **B.** Marker-panel staging scores. The first three panels show component scores for early nc14 zygotic, late nc14 zygotic, and maternally deposited transcripts; the fourth shows the staging index (late minus early), on which the statistical test is defined. Note that higher values indicate a later developmental position for the late zygotic panel and the staging index, but an earlier position for the early zygotic and maternal panels. In both panels, points are individual libraries, coloured by genetic background and shaped by sequencing batch; crossbars indicate group means and error bars ± SEM. All values are adjusted for the additive sequencing-batch effect. *P* values are from Welch’s two-sample *t*-test comparing genetic backgrounds within each temperature condition; ns, not significant. Samples of both backgrounds spanned early and late nc14, reflecting sub-stage heterogeneity within hand-selected stage 5 collections; this heterogeneity was comparable between backgrounds and renders the comparison conservative.

**Fig. S4.**
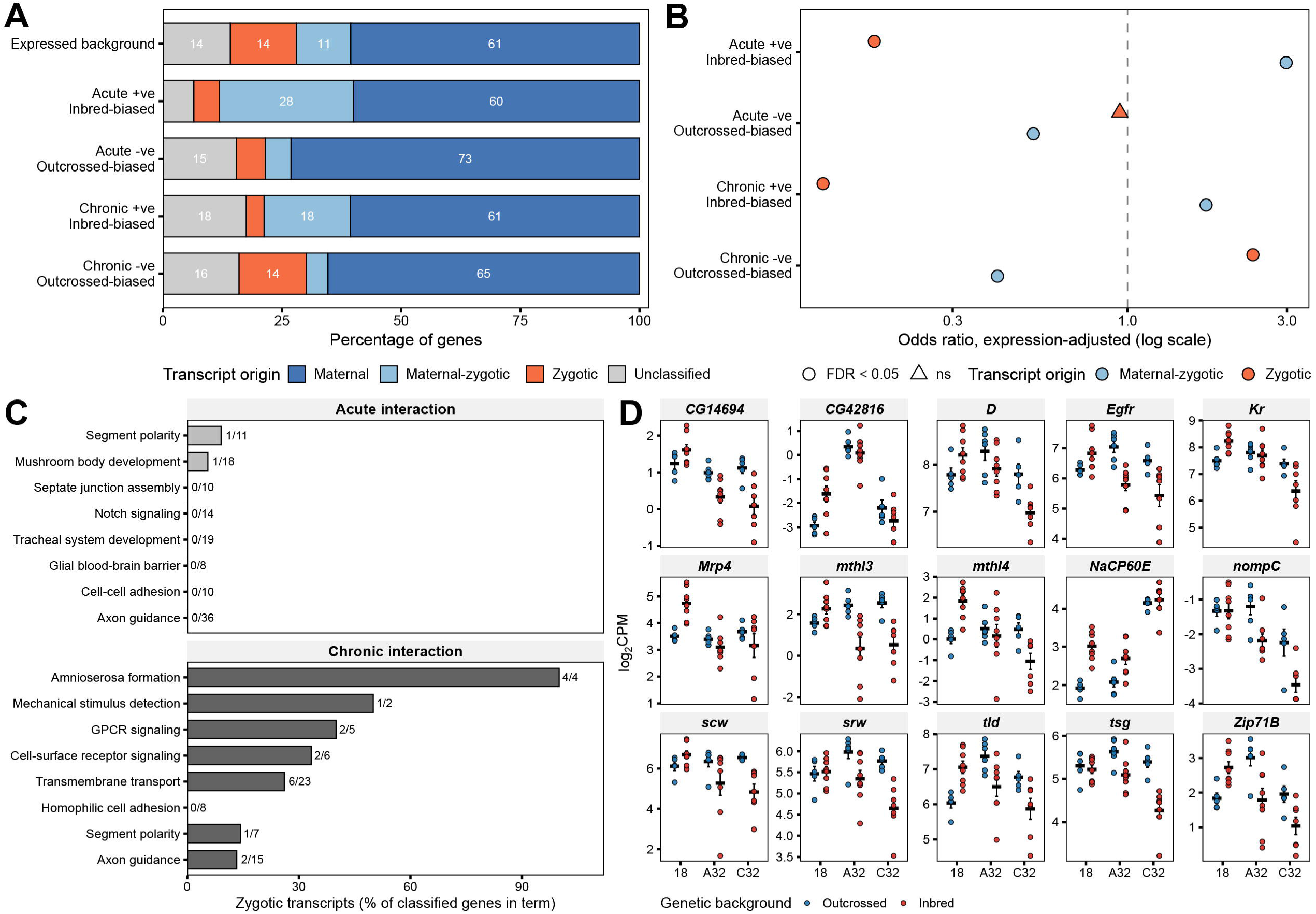
Inbred- and outcrossed-biased heat responses draw on distinct transcript pools. **A.** Composition of interaction gene sets by transcript origin, compared with the expressed background. **B.** Expression-adjusted odds ratios for enrichment of maternal-zygotic and zygotic transcripts relative to the expressed background; symbols indicate Benjamini–Hochberg significance; dashed line, no enrichment. **C.** Percentage of classified genes that are purely zygotic for each Gene Ontology term displayed in Figs. 3 and 4 for the outcrossed-biased sets; fractions indicate zygotic genes over classified genes per term. **D.** Expression of the zygotic genes contributing to chronic outcrossed-biased terms across all six sample groups. Points are individual libraries, crossbars group means, error bars ± SEM.

